# White matter microstructural abnormality precedes cortical volumetric decline in Alzheimer’s disease: evidence from data-driven disease progression modelling

**DOI:** 10.1101/2022.07.12.499784

**Authors:** CS Parker, PSJ Weston, H Zhang, NP Oxtoby, the Alzheimer’s Disease Neuroimaging Initiative

## Abstract

Sequencing the regional progression of neurodegeneration in Alzheimer’s disease (AD) informs disease mechanisms and facilitates identification and staging of individuals at greatest risk of imminent cognitive decline, which may aid the development of early therapeutic interventions. Previous attempts to sequence neurodegeneration have analysed measures of regional volume and identified the initial sites of atrophy. However, focal microstructural alterations in white matter have also been reported in early AD. Yet, the temporal ordering of abnormality in measures of white matter microstructure relative to grey matter volume has not been established. In this study we used event-based modelling of disease progression (EBM) to provide a data-driven evaluation of the temporal sequence of abnormality in markers of white matter microstructure relative to grey matter volume. Regional microstructural metrics derived from diffusion tensor imaging (DTI) and regional volumes from Freesurfer cortical parcellation were obtained from the Alzheimer’s disease Neuroimaging Initiative (ADNI) database for 441 amyloid-positive participants (81 AD-dementia, 159 mild cognitive impairment, 201 cognitively normal). The estimated sequence shows a series of abnormalities in markers of white matter microstructure, followed by sequential grey matter volumetric decline, with no overlap between the two. Analysis of positional variance and cross-validation supports the robustness of our findings. These results provide the first data-driven evidence that markers of white matter microstructural degeneration precede those of cortical volumetric decline in the AD cascade. This prompts a re-evaluation of the view that regional volumetric decline can be used to characterise the very earliest stages of AD neurodegeneration. Instead, we suggest that white matter microstructural markers provide an earlier window into AD neurodegeneration. An early staging system of AD neurodegeneration based on measures of brain microstructure may find application in selecting AD subjects with early but minimal brain damage for clinical trials that aim to prevent cognitive decline.

## 1. Introduction

Alzheimer’s disease (AD) is a progressive neurological disorder characterised by a cascade of brain abnormalities, including accumulation of amyloid-beta and hyperphosphorylated tau proteins, neuronal dysfunction, and neurodegeneration. However, while amyloid deposition occurs early, it is not closely related, either temporally or topographically, to downstream cognitive decline. Neurodegeneration, which encompasses brain structural damage at the micro- and macroscopic scale, occurs in both white and grey matter, and is the major driver of cognitive decline in AD (Englund 1998, Jack Jr. et al 1997, Frisoni et al 2002, McDonald et al 2012). Establishing the temporal evolution of AD neurodegeneration may aid the development of therapeutic interventions that target early phases of neurodegeneration prior to significant cognitive decline and irreversible neuronal loss.

Hypothetical (Jack Jr. et al 2010) and data-driven (Fonteijn et al 2012, Young et al 2014, Oxtoby & Alexander 2017) approaches have been previously used to investigate the progression of AD neurodegeneration. These investigations have established that volumetric decline occurs initially in medial temporal lobe grey matter, following tau protein deposition, and is closely correlated to cognitive decline. On the regional level, abnormality is evident in medial temporal lobe areas first (hippocampus and entorhinal cortex), then in lateral temporal lobe areas (mid-temporal gyrus and fusiform gyrus) and finally in frontal lobe areas (Jack Jr. et al 2010, Young et al 2014).

While informative on AD neurodegeneration, these approaches have relied on evidence of overt atrophy derived from measurements of regional volume. The relatively coarse description of macroscopic structural breakdown provided by regional volumetric measures does not account for microscopic neurodegenerative processes, such as alterations in neuronal morphology, organisation, and density (Englund 1998), which may be occurring in brain regions prior to measurable volumetric reductions.

Previous studies have shown that measures of brain microstructure derived from diffusion tensor imaging (DTI) (Basser et al 1994) are abnormal in white matter regions in prodromal and early phases of AD (Sexton et al 2011, Douaud 2013, Yu et al 2017, Fellgiebel et al 2018), suggesting such measures are sensitive to early AD neurodegeneration. However, the temporal progression of regional white matter microstructural abnormality relative to grey matter volumetric decline has not been established.

In this study, we use event-based modelling of disease progression (EBM) to provide a data-driven evaluation of the temporal ordering of abnormality in markers of white matter microstructure and grey matter volume in AD.

## 2. Materials and Methods

### 2.1. Overview

We leverage the recent availability of a large repository of regional measures of white matter microstructure and grey matter volume available from the Alzheimer’s disease Neuroimaging initiative (ADNI) (Mueller et al 2005). This ADNI data source and the acquisition and pre-processing steps used for its generation are described in the *ADNI imaging data section*. Selection of biomarkers and processing of ADNI data is described in the *Biomarkers of regional microstructure and volume* section. Sequencing of biomarkers is detailed in the *Event-based modelling of disease progression* section. The *Cross-validation* section describes the resampling analysis used to determine the data-dependent variability in the estimated sequence.

### 2.2. ADNI Imaging Data

#### 2.2.1. Biomarker sources

To enable sequencing of regional neurodegeneration, we required a large selection of regional biomarkers in subjects that span the entire course of AD. To obtain these, we downloaded imaging biomarkers from ADNI (Mueller et al 2005) stages 1, GO, 2 and 3 via the LONI database (adni.loni.usc.edu). ADNI was launched in 2003 by the National Institute on Aging (NIA), the National Institute of Biomedical Imaging and Bioengineering (NIBIB), the Food and Drug Administration (FDA), private pharmaceutical companies and non-profit organizations, as a $60 million, 5-year public-private partnership. For up-to-date information, see http://www.adni-info.org. Consent was obtained according to the Declaration of Helsinki and was approved by the ethical committee of the institutions in which the work was performed.

Summary spreadsheets of regional white matter microstructure derived from DTI and regional grey matter volume derived from Freesurfer were downloaded from LONI on 5^th^ June 2022 (ADNI_DTIROI_10_03_19.csv, ADNI_DTIROI_MEAN_V2.csv, and ADNIMERGE.csv). All spreadsheets were merged based on matching the subject identifiers (‘RID’ column) and visit codes (‘VISCODE’ column).

The image acquisition and processing steps undertaken by ADNI to generate the data for the summary spreadsheets are described below.

#### 2.2.2. Image acquisition

Imaging data were derived from DTI and T1-weighted images acquired on a 3T GE scanner using standard ADNI protocols, as described previously (Jack et al 2008, Nir et al 2013). Subjects were scanned at multiple sites across North America. T1-weighted imaging consisted of a spoiled gradient echo sequence with 256×256 matrix, voxel size 1.2×1.0×1.0mm3, TI=400ms, TR=6.98ms, TE=2.85ms and flip angle =11°. DTI consisted of a spin-echo echo planar imaging (EPI) sequence with 256×256 matrix, voxel size 2.7×2.7×2.7mm3 and TR=9000ms. Each DTI contained 5 images with no diffusion weighting and 41 diffusion-weighted images with b=1000 s/mm^2^, with a total scan time of 9 mins. The quality of T1-weighted and DTI images were checked visually and those with excessive motion or image artefacts were excluded.

#### 2.2.2. Image pre-processing

Following acquisition, subjects’ DTI data were brain-masked and corrected for motion and eddy-current induced distortions using FSL’s eddy_correct tool (www.fmrib.ox.ac.uk/fsl). Phase-susceptibility induced distortions were corrected by aligning the first non-diffusion-weighted volume to the respective subjects’ T1-weighted image using FSL flirt followed by elastic registration using inverse consistent registration algorithm (Leow 2007). Diffusion tensors were fitted using FSL dtifit and voxel-wise diffusion tensor parameters of fractional anisotropy (FA), mean diffusivity (MD), axial diffusivity (AxD) and radial diffusivity (RD) were derived from the tensor eigen-values. The JHU atlas (Mori et al 2008) FA image was aligned to each subjects’ FA image and the corresponding deformation applied to the Eve white matter atlas to map it to each subjects native space (Nir et al 2013). The arithmetic mean of each DTI metric was then calculated in fifty-two regions-of-interest (ROIs) defined in the Eve white matter atlas. Images with processing failures were excluded.

Extra-cerebral tissue was removed from T1-weighted images using the robust automated brain extraction (ROBEX) (Iglesius et al 2011) and intensity inhomogeneity normalized using the MNI nu_correct tool (www.bic.mni.mcgill.ca/software/). The brain was then extracted using FSL bet (Smith et al 2002). The cortical grey matter regional volumes were derived from Freesurfer (version 5.1) parcellation (Fischl et al 2004). T1-weighted images with processing failures were also excluded.

### 2.3. Biomarkers of regional microstructure and volume

#### 2.3.1. Normalization

Regional white matter microstructure biomarkers derived from DTI were averaged over left and right hemispheres, as we did not expect any hemisphere-specific disease effects. Grey matter volumes were also averaged over hemispheres and normalised by intra-cranial volume to correct for individual differences in head size (Marinescu et al 2021).

#### 2.3.2. Biomarker selection

Ten white matter ROIs showing significant microstructural abnormality at the group level in previous studies of AD were selected for disease progression modelling (Rose et al 2000, Duan et al 2006, Yasmin et al 2008, Fellgiebel et al 2008, Douaud et al 2011, Oishi er al 2012, Kheihaninijad et al 2013) (Table 1). To provide sensitivity to the initial microscopic degenerative process occurring in each region, the DTI metric with earliest abnormality was selected for each ROI. In the absence of an established method for determining the earliest abnormality, the DTI metric in the earliest sequence position following an ROI- and DTI-specific EBM was chosen. The DTI metric (FA, MD, RD or AxD) used for each ROI in the final EBM is shown in Table 1, column 3.

**Table 1.** List of region names, abbreviations and biomarkers for EBM.

| Region |  | Biomarker |  |  |  |  |
| --- | --- | --- | --- | --- | --- | --- |
| Abbreviation | Name | WM microstructure |  |  |  | GM volume |
|  |  | FA | AxD | RD | MD | Volume |
| *White matter* |  |  |  |  |  |  |
| BCC | Body of corpus callosum |  | ✓ |  |  |  |
| CGC | Cingulum |  |  |  | ✓ |  |
| CGH | Hippocampal cingulum |  | ✓ |  |  |  |
| FX | Fornix body |  |  |  | ✓ |  |
| FX-ST | Fornix limbs or stria terminalis |  | ✓ |  |  |  |
| GCC | Genu of corpus callosum | ✓ |  |  |  |  |
| SCC | Splenium of corpus callosum |  | ✓ |  |  |  |
| SLF | Superior longitudinal fasciculus |  |  | ✓ |  |  |
| SS | Sagittal stratum |  |  |  | ✓ |  |
| UNC | Uncinate fasciculus | ✓ |  |  |  |  |
| *Grey matter* |  |  |  |  |  |  |
| HIP | Hippocampus |  |  |  |  | ✓ |
| ENT | Entorhinal cortex |  |  |  |  | ✓ |
| MID | Mid-temporal gyrus |  |  |  |  | ✓ |
| FUS | Fusiform gyrus |  |  |  |  | ✓ |

The four grey matter ROIs selected as reference points for known macroscopic degeneration in AD were the hippocampus, entorhinal cortex, fusiform gyrus and mid-temporal gyrus. These regions are recognised as the earliest sites of volumetric atrophy in typical AD (Jack Jr. et al 1997, 2010, Young et al 2014).

The abbreviations, region names and metrics used in the final EBM are listed in Table 1.

#### 2.3.2. Subjects

We included data from subjects with all available biomarkers, who tested positive for brain amyloid-beta plaques following cerebrospinal fluid (CSF) or positron emission tomography (PET) testing. This yielded data for 441 AD subjects: 89 AD-dementia patients, 159 individuals with mild cognitive impairment (MCI) or subjective memory complaints, and 201 cognitively normal subjects. Note that in the case where multiple timepoints were available with all biomarker data the earliest timepoint was chosen. Subject demographics are shown in Table 2.

**Table 2.** Demographics of AD subjects with available data for all biomarkers. The threshold for amyloid-positivity for CSF amyloid-beta was < 992 pg/ml and for PET amyloid-beta positivity was SUVR > 1.1 for Pittsburgh compound B, Florbetapir or Florbetaben radiotracers.

| Demographics | Cognitively normal | MCI | AD-dementia |
| --- | --- | --- | --- |
| n | 201 | 159 | 89 |
| Sex M/F | 81/120 (40%) | 91/68 (57%) | 47/34 (58%) |
| Age (yrs ± s.d.) | 72 ± 6 | 73 ± 7 | 73 ± 8 |
| Education (yrs ± s.d.) | 16.7 ± 2.3 | 15.8 ± 2.7 | 15.7 ± 2.6 |
| APOE4 +/- | 93/104 (31%) | 95/58 (62%) | 55/24 (70%) |

### 2.4. Event-based modelling of disease progression

#### 2.4.1. Event sequence

EBM represents the evolution of biomarker abnormalities as a sequence of events, S, which is a permutation of the biomarker indices (Fonteijn et al 2012). The sequence describes the order of events, each signifying that a biomarker has transitioned from a normal state (¬E) to abnormal state (E).

The estimated sequence of events is determined by maximum likelihood estimation, with the likelihood being the probability of observing the biomarker data, X, given a particular sequence, S. Assuming (i) an equal probability of each stage, k, in the sequence, (ii) independence of subjects’ data, and (iii) independence of the different biomarker observations at each sequence stage, the likelihood of the sequence is:

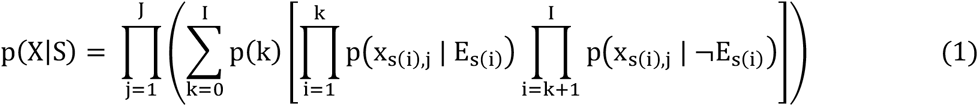

where J is the number of subjects, I is the number of biomarkers and the subscript S(i) is the biomarker index at the i’th position in the sequence. The stage of each individual subject is considered unknown and is therefore marginalised out.

The sequence with maximum likelihood was determined using 50 stochastic greedy ascent realisations (1000 iterations each), each initialised with a random sequence. For each iteration two events were chosen at random, their positions swapped, and the likelihood of the sequence evaluated using Eq. 1. The estimated sequence was that with highest likelihood over all greedy ascent initialisations and iterations.

#### 2.4.2. Event distributions

Evaluating the sequence likelihood requires the likelihood functions of each biomarkers’ data under the condition that the event has or has not occurred, p(X|E) and p(X|¬E). To estimate these distributions, we fitted a kernel density mixture model to each biomarkers data (Firth et al 2020, https://github.com/ucl-pond/kde_ebm). As kernel density estimation is a non-parametric method for estimating probability distributions, the event distributions derived for each biomarker are more robust to non-gaussian distributions. We used the subjects’ clinical diagnosis (cognitively normal or AD-dementia) to initialise the kernel density estimation (Firth et al 2020).

#### 2.4.3. Sequence uncertainty

Positional uncertainty in the sequence was quantified using Markov chain Monte Carlo (MCMC) sampling, as described in (Fonteijn et al 2012). Following initialisation at the maximum likelihood sequence, MCMC iteratively sampled 100,000 sequences and calculated their likelihoods. Using these MCMC samples, the likelihood for each biomarker event being at each position in the sequence was 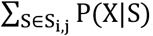, where S_i,j_ is the set of all sequences with biomarker event i at position j. The set of positional likelihoods was visualised in a positional variance diagram, which summarises positional uncertainty as a matrix where each entry (i, j) is the probability of event i being at position j (Young et al 2014).

#### 2.4.4. Cross-validation

To stress-test the dependence of the sequence on the sample, variability in the sequence was assessed using cross-validation style data resampling (as in Young et al 2014). The resampling analysis estimated the sequence from a total of 450 resamples of subjects’ data generated using a stratified (by diagnosis) repeated 9-fold cross-validation style resampling. The positional uncertainty across resamples was summarised in a positional variance diagram where each entry (i, j) is the relative frequency across resamples where biomarker event i occurred at position j (Young et al 2014).

#### 2.4.5. Data availability

ADNI data used for this study can be obtained from LONI database (adni.loni.usc.edu). Scripts for data processing are available from the KDE-EBM package, (https://github.com/ucl-pond/kde_ebm).

## 3. Results and Discussion

### 3.1. The sequence of white matter microstructural abnormality and grey matter volumetric decline in AD

The sequence of white matter microstructural and grey matter volumetric neurodegeneration in AD is shown in Fig. 1, and suggests that white matter microstructural abnormality precedes grey matter volumetric abnormality. Specifically, the sequence of biomarker abnormalities proceeds as follows: 1. increased MD in the body of the fornix, 2. increased AxD in the body of the corpus callosum, 3. increased AxD in the fornix striaterminalis, 4. increased MD in the sagittal stratum, 5. increased AxD in the splenium of the corpus callosum, 6. increased MD in the cingulate gyrus, 7. increased RD in the superior longitudinal fasciculus, 8. decreased FA in the genu of the corpus callosum, 9. increased AxD in the hippocampal cingulum, 10. decreased FA in the uncinate fasciculus, 11. decreased volume of the entorhinal cortex, 12. decreased volume of the fusiform gyrus, 13. decreased volume of the mid-temporal gyrus and 14. decreased volume of the hippocampus.

**Figure 1.**
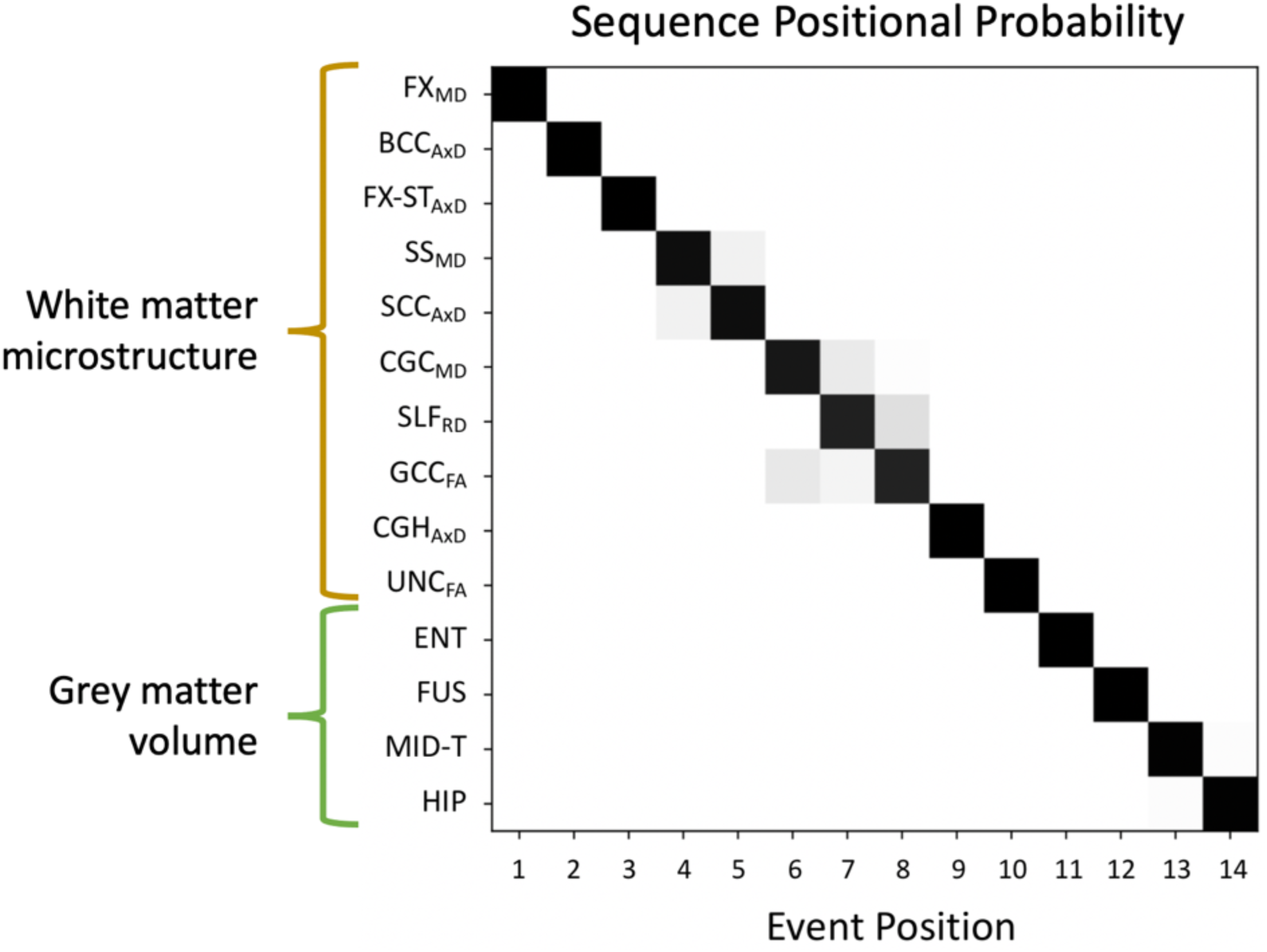
Positional variance diagram showing the estimated sequence (row order, top to bottom) and the uncertainty associated with each biomarkers’ event position (row, column entries). Each entry P_i,j_ of the diagram shows the relative probabilities of biomarker i (row) being at position j (column) in the sequence.

### 3.2. Sequence positional uncertainty

Analysis of sequence uncertainty using MCMC shows that positional variance is low, suggesting high confidence in the ordering (Fig. 1).

Cross-validation shows that the estimated sequence is consistent (Fig. 2), with the median position of each biomarker event being equal to its position estimated on the original dataset. Importantly, the relative position of markers of white matter microstructure and markers of grey matter volume was preserved, supporting the generalisability of our findings that WM microstructural abnormality precedes GM volume abnormality.

**Figure 2.**
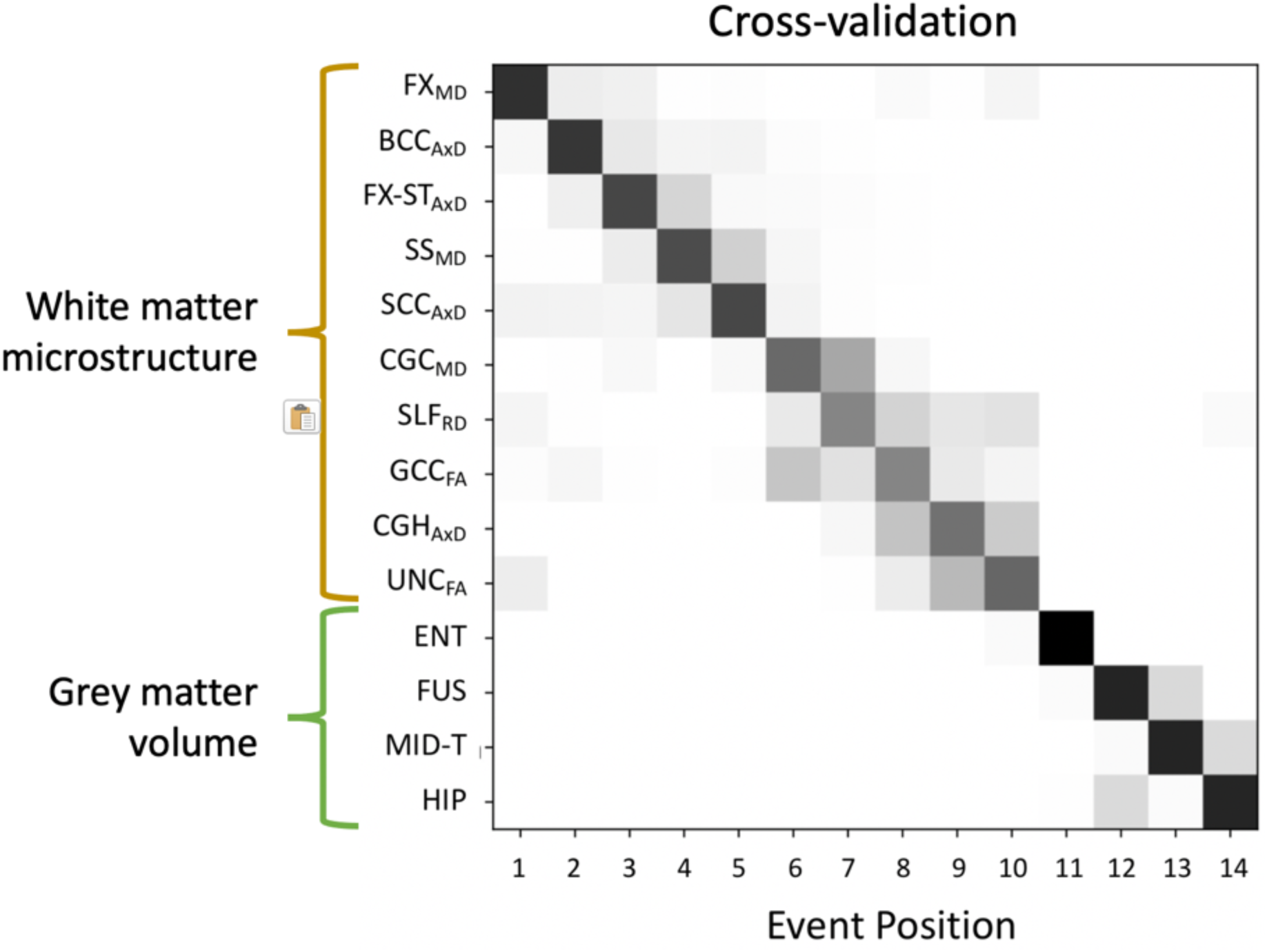
Positional variance diagram showing the relative frequency of each biomarker position across the set of sequences generated using a cross-validation style resampling.

The entorhinal cortex consistently separated white matter microstructural events from later grey matter volumetric events. Among the grey matter regions, the early positioning of the entorhinal cortex is consistent with post-mortem Braak staging which shows that entorhinal cortex is the earliest site of tau deposition and one of the earliest grey matter regions to undergo neurodegeneration in AD (Braak, H & Braak, E 1991).

A moderate positional variation was observed across resamples within groups of markers of white matter microstructure and grey matter. For example, increased MD in the fornix body was sometimes positioned at later stages while decreased FA in the uncinate fasciculus was sometimes positioned at earlier stages. Furthermore, reduced volume of the hippocampus was frequently positioned immediately following entorhinal cortex volume reductions, which is consistent with it being recognised as one of the preliminary sites of grey matter volumetric decline in AD (Jack Jr. et al 1997). The sample-dependent positional variation may be due to the presence of sub-types of AD, with differing spatial patterns of spread, within the original dataset. Future work can explore the possibility of AD neurodegeneration sub-types (Young et al 2018, Vogel et al 2021).

### 3.3. Sequencing white matter microstructural neurodegeneration in AD: novelty, mechanisms and applications

Previous studies have demonstrated white matter microstructural abnormality in early stages of AD by analysing group differences (Sexton et al 2011, Yu et al 2017). However, none have so far determined their positioning in the AD cascade relative to grey matter volumetric decline. Our study uses a data-driven approach to show that markers of white matter microstructure become abnormal earlier in the AD cascade than grey matter volumetric decline. This finding suggests that methods to stage AD based on regional grey matter volumes may be omitting earlier microscopic neurodegenerative processes occurring in white matter that is useful for pre-symptomatic staging. A staging system for early AD neurodegeneration should therefore include markers of white matter microstructure.

Multiple mechanisms have been proposed to explain white matter microstructural neurodegeneration in AD. Wallerian degeneration postulates that white matter tracts connected to degenerating grey matter regions undergo degeneration (Beaulieu et al 1996, Pierpaoli et al 2001). The retrogenesis hypothesis suggests that late-myelinating white matter structures are the earliest to deteriorate in AD (Reisberg et al 1999), followed by earlier-myelinating structures. Neuroinflammation and spread of tau may also contribute to white matter degeneration (Akiyama et al 2000, Powell et al 2018). To provide insight into the relative contributions of these mechanisms, future studies will explore the positioning in markers of grey matter microstructure, which were not included in this analysis due to the confounding effect of CSF partial volume. Mechanistic modelling of disease spread (Raj et al 2018) may also inform the extent to which each potential mechanism of white matter neurodegeneration explains the observed sequence of neurodegeneration.

Early staging of neurodegeneration has potential application for advancing preventative treatment development, monitoring individual progression, and predicting future disease progression. By identifying subjects at the disease stage where cognitive is immanent due to early neurodegeneration, individuals can be identified in whom cognitive decline can be prevented and in whom treatment effects based on cognitive endpoints, which are frequently used in early AD trials (Andrieu et al 2015), are detectable in short-term studies. Furthermore, stage-based selection can identify individuals with similar underlying biochemistry and disease who are likely to respond consistently to a single treatment, thus improving treatment detectability (Oxtoby & Alexander 2017). Staging markers of regional white matter microstructure has the additional benefit of allowing fine-grained staging, which may allow more detailed monitoring and prognosis of individual disease progression. Future work will stage subjects within the sequence and characterise the cross-sectional and longitudinal AD cognitive profiles of each stage to determine the optimal stage to target for clinical trials.

In comparison to PET imaging of tau accumulation, which has also been associated with cognitive decline, staging of neurodegeneration has several advantages. Firstly, cognitive decline is more closely correlated to neurodegeneration than tau accumulation (Jack Jr. et al 2010). This suggests a closer association between neurodegeneration-based stages and cognitive decline, and therefore more accurate predictions of cognitive outcomes and expected preventative treatment effects. Secondly, microstructural imaging uses diffusion-weighted magnetic resonance imaging, which is less expensive than PET imaging and is also non-invasive, which makes it easier to adopt in clinical settings.

### 3.4. An updated hypothetical model of Alzheimer’s disease

Based on the observed sequence, we provide an updated hypotheses on the temporal evolution of brain structural neurodegeneration in AD, where abnormalities in markers of white matter microstructure precede measures of volumetric decline. This hypothesis is visualised as an updated version of the AD pathophysiological cascade introduced by (Jack Jr. et al 2010) (Fig. 3). The early positioning of microstructural abnormality complements the hypothesis of (Weston et al 2015), who used systematic review to suggest that microstructural changes in grey matter precedes volumetric decline. In future, joint analysis of white and grey matter changes in microstructure and volume are needed to sequence the evolution of microscopic and macroscopic neurodegeneration in AD.

**Figure 3.**
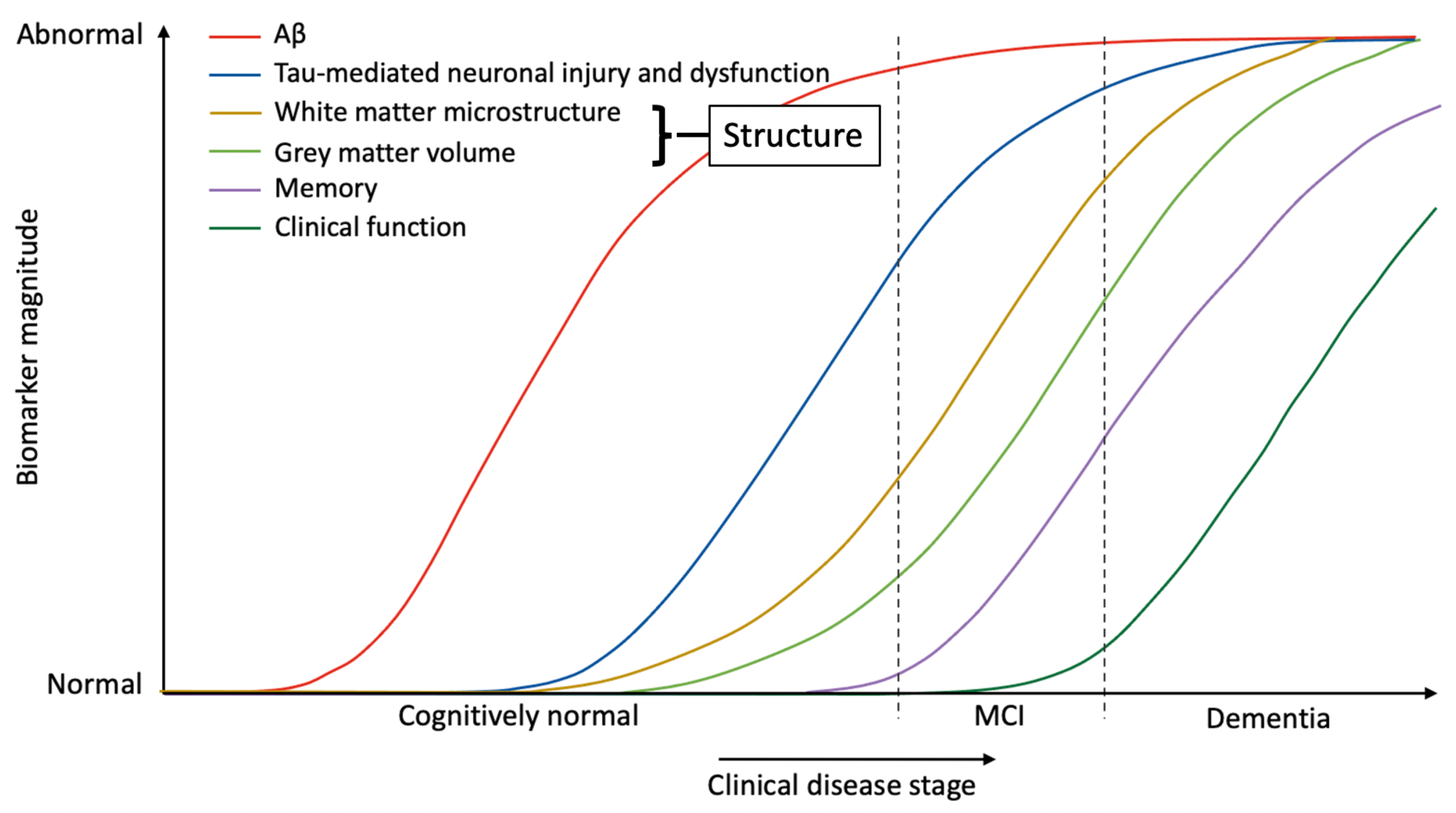
Updated model of the temporal sequence of biomarker abnormalities in AD (adapted from the model proposed by Jack Jr. et al 2010). In the original model (Jack Jr. et al 2010), structural brain degeneration was considered a single trajectory and referred primarily to measures of grey matter volume. We propose to split structural brain degeneration into earlier white matter microstructural abnormality and later grey matter volumetric decline.

## 4. Conclusion

Our study provides the first data-driven evidence that markers of white matter microstructure become abnormal prior to those of grey matter volume in the AD neurodegenerative cascade. This prompts a re-evaluation of the view that the earliest brain structural breakdown in AD can be characterised using volumetric measures. Indeed, our results suggest that measures of microstructure are essential for investigation of the earliest neurodegeneration in AD. A staging system based on white matter microstructural abnormality may be used to select homogeneous groups of early-stage AD subjects for participation in clinical trials that aim to ameliorate or prevent the earliest signs of cognitive decline, and can also enable more precise, fine-grained tracking and prognosis of progressive neurodegeneration over time.

## Acknowledgments

CSP, DCA and HZ are supported by the Medical Research Council (MR/T046473/1). CSP is further supported by the EPSRC CMIC Platform Grant (EP/M020533/1). NPO is a UKRI Future Leaders Fellow (MR/S03546X/1) and acknowledges funding from the E-DADS project (EU JPND 2019; UK MRC MR/T046422/1), and the National Institute for Health Research University College London Hospitals Biomedical Research Centre.

